# Identifying suitable *Listeria innocua* strains as surrogates for *Listeria monocytogenes* for horticultural products

**DOI:** 10.1101/586016

**Authors:** Vathsala Mohan, Reginald Wibisono, Lana de Hoop, Graeme Summers, Graham C Fletcher

**Author notes:** Current address: MediaCom, London, UK.

## Abstract

We conducted a laboratory-based study testing nine *Listeria innocua* strains independently and a cocktail of 11 *Listeria monocytogenes* strains. The aim was to identify suitable *L. innocua* strain(s) to model *L. monocytogenes* in inactivation experiments. Three separate inactivation procedures and a hurdle combination of the three were employed: thermal inactivation (55°C), UV-C irradiation (245 nm) and chemical sanitiser (Tsunami™ 100, a mixture of acetic acid, peroxyacetic acid and hydrogen peroxide). The responses were strain dependent in the case of *L. innocua* with different strains responding differently to different regimes. *L. innocua* isolates generally responded differently to the *L. monocytogenes* cocktail and had different responses among themselves. In the thermal inactivation treatment, inactivation of all strains including the *L. monocytogenes* cocktail plateaued after 120 minutes. Chemical sanitiser, inactivation could be achieved at concentrations of 10 and 20 ppm with inactivation increasing with contact time up to 8 minutes, beyond which there was no significant benefit. Although most of the *L. innocua* strains in the study responded similarly to *L. monocytogenes* when subjected to a single inactivation treatment, when the treatments were applied as hurdle, all *L. innocua* strains except PFR16D08 were more sensitive than the *L. monocytogenes* cocktail. PFR16D08 almost matched the resistance of the *L. monocytogenes* cocktail but was much more resistant to the individual treaments. A cocktail of two *L. innocua* strains (PFR 05A07 and PFR 05A10) had the closest responses to the hurdle treatment to those of the *L. monocytogenes* cocktail and is therefore recommended for hurdle experiments.

**Importance:** Owing to researcher safety risks it is often difficult to use actual pathogens, such as *Listeria monocytogenes*, to explore different inactivation procedures under field conditions. Organisms that are closely related to the pathogen but without its virulence are therefore used as surrogates for the actual pathogen. However, this assumes that the surrogate will behave in a similar manner to the pathogen and it is difficult to predict the responses of the surrogate compared to the actual pathogen. This study compares the responses of individual and combined “cocktails” of strains of non-pathogenic *Listeria innocua* to different inactivation procedures when compared to the response of a cocktail of *L. monocytogenes*. Our study highlights the importance of evaluating a number of strains when choosing surrogates.

## Introduction

*Listeria monocytogenes* is a Gram-positive facultative anaerobe that is found in a range of natural environments (including soil, water and vegetation); in food-processing environments and in ready-to-eat food products. Ingestion of *L. monocytogenes* can cause serious illness in pregnant women, neonates, elderly and immune-compromised individuals (1). Its ability to grow at a broad range of temperatures from −7°C to 45°C (2), in salt at concentrations up to 10%, and a pH range from 4.1 to 9.6 makes *L. monocytogenes* a very significant and robust foodborne pathogen (3).

Control measures including physical and chemical treatments have greatly reduced the prevalence of *L. monocytogenes* in a variety of food products and in food-processing environments. (4, 5). However, the human disease incidence rate has not decreased over past decades (3, 6). Incidence of human listeriosis cases caused by *L. monocytogenes* averages at around one per 200,000 people in New Zealand, with an estimated 84.9% of cases being food related (7). Although regulatory controls and industry actions have been in place for many years in the US, listeriosis outbreaks from dairy products showed no decrease in frequency (8), and outbreaks from fresh horticultural products have been a concern in the past decade (8–11). Fresh produce is often eaten raw so the high temperatures that are used to eliminate pathogens from other food products cannot be used, meaning that other control strategies must be found.

Because of the pathogenicity and the environmentally persistent nature of *L. monocytogenes*, it is challenging to safely conduct large-scale experiments using *L. monocytogenes* in research pilot plants or commercial settings. A surrogate bacterium is often sought that has similar genotypic as well as phenotypic characteristics as using surrogates gives a safety margin to protect researchers by preventing exposure to pathogens (12–14). *L. monocytogenes* and *L. innocua* are genetically similar and until 1981 the two were not recognised as separate species (15). Since then, comparative genomic studies have differentiated hundreds of strain-specific genes for these two bacterial species (16).

*L. innocua* is a non-pathogenic *Listeria* spp. found in similar environments to *L. monocytogenes*. The main phenotypic characteristic that distinguishes it from *L. monocytogenes* is that it is not haemolytic (17–19). Enhanced haemolytic activity testing, also known as the Christie, Atkins, Munch-Petersen (CAMP) test has been employed regularly for differentiating *L. innocua* from *L. monocytogenes* (20–22). However, some strains of *L. monocytogenes* have now been shown to be non-haemolytic (23). Apart from some studies on heat inactivation in milk and meat (24–26), there have been no studies on selecting suitable strains of *L. innocua* as surrogate organisms to investigate non-thermal inactivation procedures. Although strains of *L. innocua* have been used as surrogates (27), there are gaps in previous research and one gap is that no one study has yet tested that *L. monocytogenes* and *L. innocua* behave similarly using non-thermal inactivation. Regarding previous thermal inactivation procedures, a study in hamburger patties identified *L. innocua* strains M1, and SLCC5640 to be good thermal processing surrogate models for *L. monocytogenes*. Furthermore, in that study, *L. innocua* M1 was identified as the preferred surrogate as it is more thermo-tolerant than *L. monocytogenes* and can provide a margin of safety in the evaluation for effectiveness of heat treatments for *L. monocytogenes* (25). A study of treatments for fresh-cut lettuce found that a mild heat treatment could actually increase the growth of *L. monocytogenes* during storage (28). Other non-thermal inactivation procedures include using ultraviolet irradiation (UV) and sanitisers. UV irradiation as a means of disinfection of food products is considered to be cheap and clean, that is it leaves no chemical residues. It can be used in combination with other disinfection processes to ensure the safety of products (29–31). UV is an electromagnetic radiation that has a wavelength ranging from 100–400 nm, shorter than that of visible light (400–700 nm), while longer than x-rays (<100 nm) (32). UV-C irradiation ranges between 200 nm to 280 nm (33). It damages bacterial and viral genetic material (29). The UV-C spectrum range of 250– 270 nm is strongly absorbed by microorganisms and is considered to be the most lethal range of wavelengths, with 262 nm being the peak germicidal wavelength (30). J. M. Oteiza et al. (34) found that UV exposure was an efficient method to treat fruit juice contaminated with *Escherichia coli* O157:H7.

C. H. Sommers et al. (35) investigated *L. monocytogenes* and *L. innocua* using UV-C at 254 nm and found that a dose of 4 J/cm2 achieved a 0.37 log_10_ reduction in *L. innocua* and a 1.93 log_10_ reduction in *L. monocytogenes* on frankfurters. They attributed the difference to the surface topography of the meats and the ingredients of the frankfurters. Similarly, sanitisers have been employed as a common method for disinfection, particularly of processing environments, equipment and wash waters. One such post-harvest sanitiser is Tsunami™ 100 (Ecolab Inc., Minnesota, USA), a mixture of acetic acid, peroxyacetic acid and hydrogen peroxide (36). It has been shown to be an effective treatment to inactivate spoilage and pathogenic bacteria including *E. coli*, *Listeria*, and *Salmonella* with its efficacy depending on the contact time, temperature, and concentration (37, 38). Yet another emerging inactivation technology for improving the quality of food and its shelf life is hurdle technology. This concept of applying a series of hurdles in the post-harvest period was developed to address consumer demands for fresh and safe produce. It is an intelligent combination of more than one post-harvest antimicrobial treatment to secure microbial safety and stability, retaining the organoleptic and nutritional quality of food products (39). and these strategies have been used for meat, fish, milk and vegetables for years (39). Some of more recent hurdle technologies include nano-thermosonication, ultrahigh pressure, photodynamic inactivation, modified atmosphere packaging of both non-respiring and respiring products, edible coatings, ethanol and products to control Maillard reactions. These have been gaining popularity in the recent years (reviewed by Gayan et al. and by Pundhir & Murtaza (40, 41)).

Considering that our main aim was to select suitable *L. innocua* surrogate (s), we wanted to employ thermal, non-thermal and a combination of treatments (hurdle) technology in the selection of surrogates. Although various strains of *L. innocua* have been used as surrogates in experimental treatments for inactivation of *L. monocytogenes*, few have been validated against *L. monocytogenes* (25). There are none that perfectly match *L. monocytogenes* and none that have been tested against the range of treatment regimes, we are interested in. The suitability of a potential surrogate varies depending on the *L. innocua* strain, food matrices, and parameters used for testing (26). Here, we report a laboratory-based study conducted on different *L. innocua* and *L. monocytogenes* strains isolated in New Zealand from different sources, in order to select suitable *L. innocua* candidates to be used as surrogates in horticultural produce using thermal, non-thermal (UV and sanitiser) treatments and the combinations (hurdle technology).

## Results

### Genotyping and selection of Listeria strains

**The *Listeria* strains were genotyped by pulse field gel electrophoresis (PFGE) to select the candidate to be used in the inactivation treatments using *Asc*I and *Apa*I enzymes. Figures 1A and 1B represent the PFGE patterns of the *L. monocytogenes* and *L. innocua*.** Nine genetically dissimilar *L. innocua*, and 11 *L. monocytogenes* strains (combined in a cocktail, referred to as *LM* cocktail for convenience throughout the manuscript) were compared by treating them with UV-C, sanitiser and heat and a combination of these three. Figure 1a shows the dendrogram of the 10 *L. innocua* strains selected (based on at least 35% genetic dissimilarity) from 19 isolates, while Figure 1b shows the dendrogram of the *L. monocytogenes* strains selected for the cocktail.

**Figure 1:**
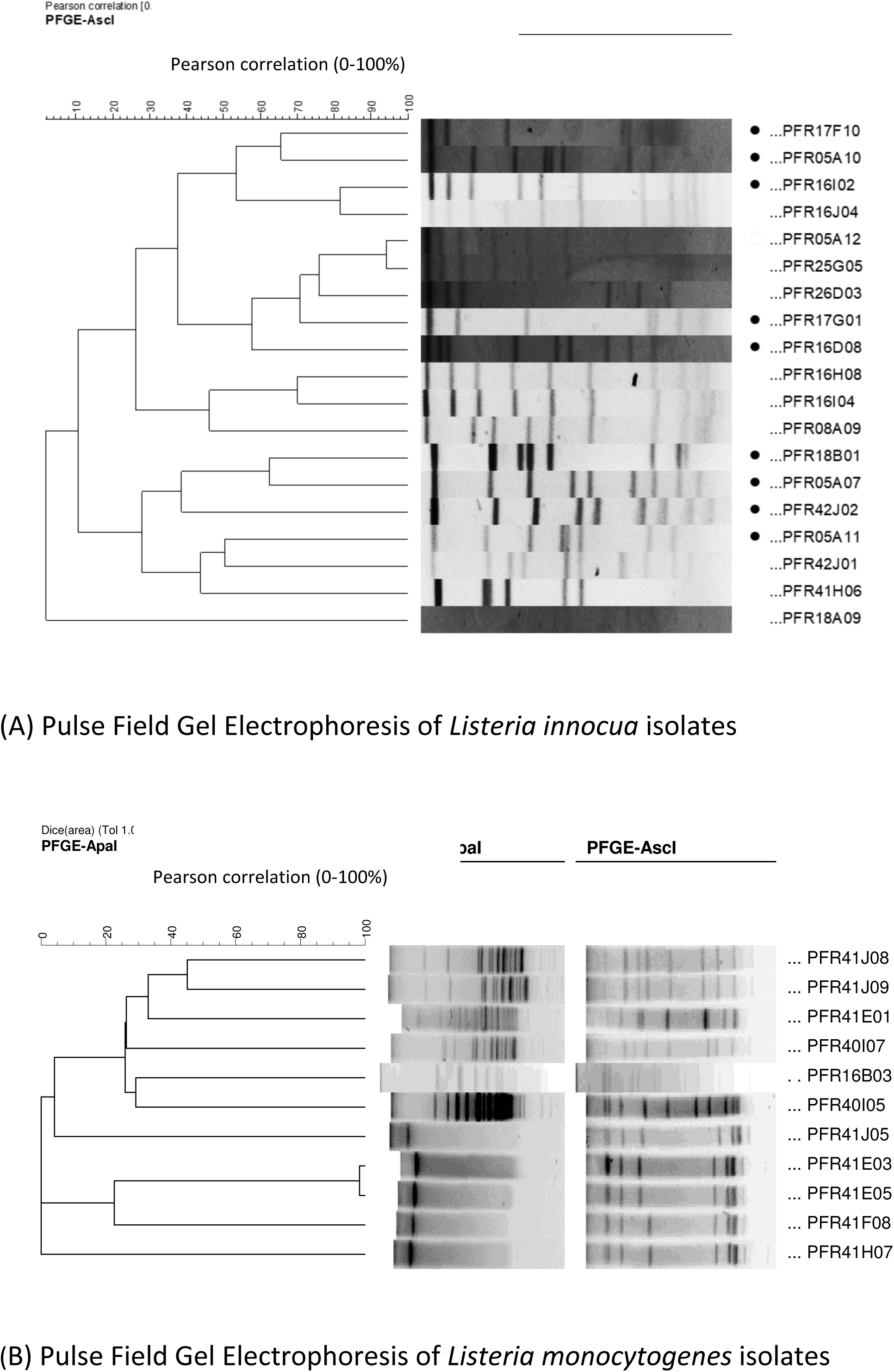
Cluster analysis of Pulse Field Gel Electrophoresis patterns performed in InfoQuest™ FP software: (A) using *Asc*I for non-pathogenic *Listeria innocua* strains. The selected isolates of *L. innocua* are indicated with black dots in front their names; (B) using *Apa*I and *Asc*I restriction enzymes for *L. monocytogenes* strains used in the *LM* cocktail.

### UV-C treatment

As a non-thermal inactivation procedure, UV-C (245 nm) was used to investigate the effect of UV-C exposure on the survivability of various strains of *L. innocua* and the *LM* cocktail. Figure 2 represents the behaviour of different strains of *L. innocua* and the LM cocktail. Increasing the dose of UV-C exposure resulted in an increase in killing effect although reductions in counts were minor for some strains (e.g. PFR 17G901 and PFR 17F10). At exposure doses of 600 and 672 mJ/cm^2^, all strains responded in a similar manner with log_10_ reductions ranging from 0.5 to 1.75. As the dose was increased to above 1000 mJ/cm^2^, different responses were observed for different strains. The *LM* cocktail had the greatest reductions in counts and only strains PFR 05A10 and PFR 18B01 had similar responses to that of the *LM* cocktail while the other strains clustered and were more resistant.

**Figure 2.**
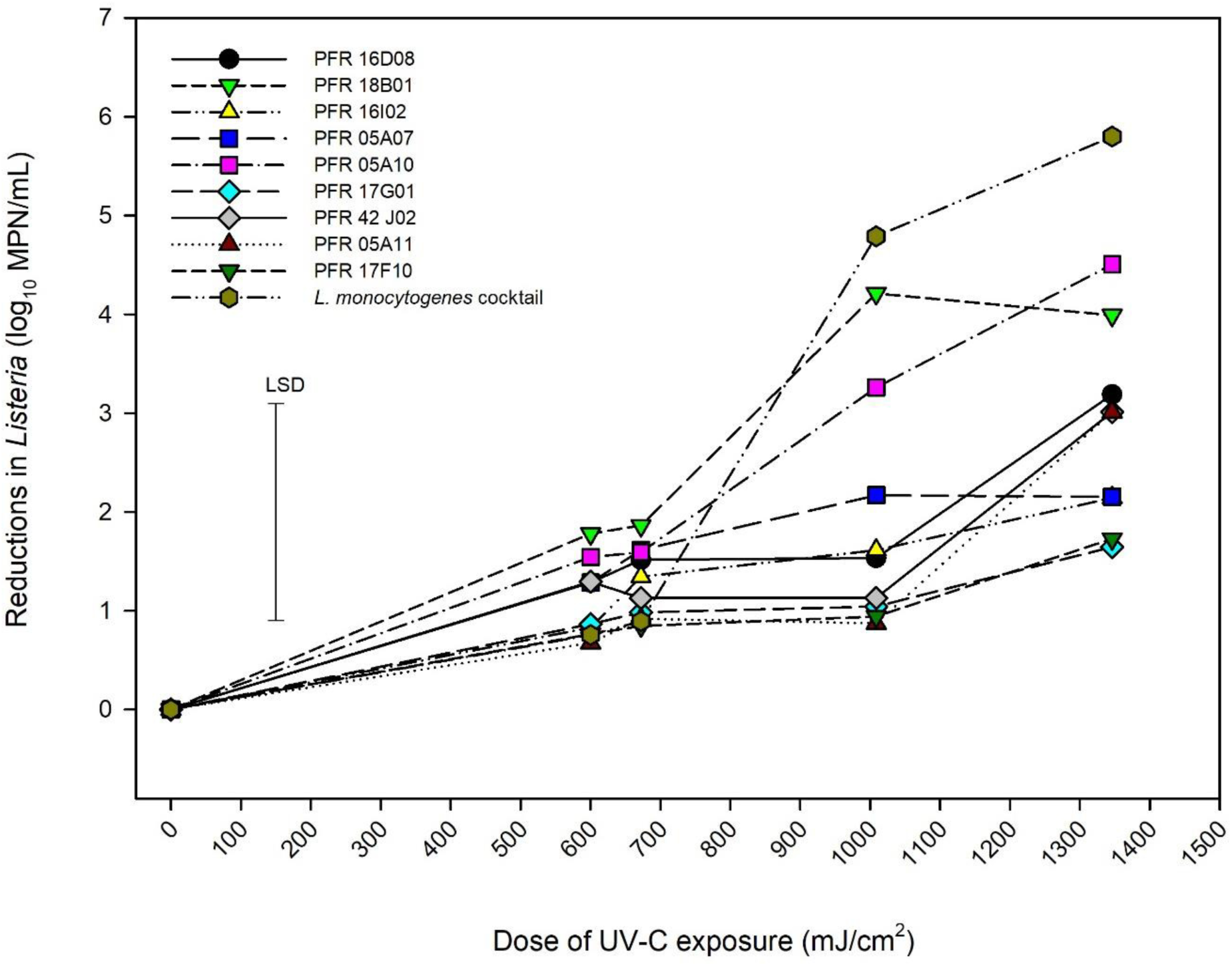
Reductions in numbers of *Listeria innocua* and of organisms in a *Listeria monocytogenes* cocktail when exposed to UV-C radiation. Error bar = least significant difference (*p* = 0.05, n = 2).

### Sanitiser treatment

Another non-thermal inactivation treatment employed was sanitiser (Tsunami™ 100). Figures 3 A, B, C and D show the pattern of reductions in *Listeria* count when treated with Tsunami™ 100 at different peroxide equivalent concentrations of 10, 20, 40 and 80 ppm, respectively. As expected, increasing contact time between the bacteria and Tsunami™ 100 led to increased kill. No reductions in counts were observed during the first 2 min of exposure to 10 ppm Tsunami™ 100 whereas reductions started to be observed during the first minute at the higher concentrations. Maximum inactivation was achieved by 8 min at concentrations of 10 and 20 ppm while this was mostly achieved within 2 min at 40 and 80 ppm. Increasing concentrations gave increasing maximum reductions in *Listeria* numbers. All strains behaved similarly to each other on exposure to 40 and 80 ppm Tsunami™ 100 while significant differences were observed between strains at 10 and 20 ppm. At 10 ppm the *LM* cocktail was more sensitive to Tsunami™ 100 than many of the *L. innocua* strains with PFR 16D08, PFR 17F10, PFR 05A07 and PFR 18B01 being significantly more resistant than *L. monocytogenes* after 8- and 16-min exposure. At 20 ppm, just 05A12 showed significantly more resistance after 4 min exposure but not at other time periods. *L. innocua strains* PFR 16I02 and PFR 17G01 always responded in a similar manner to the *LM* cocktail.

**Figure 3.**
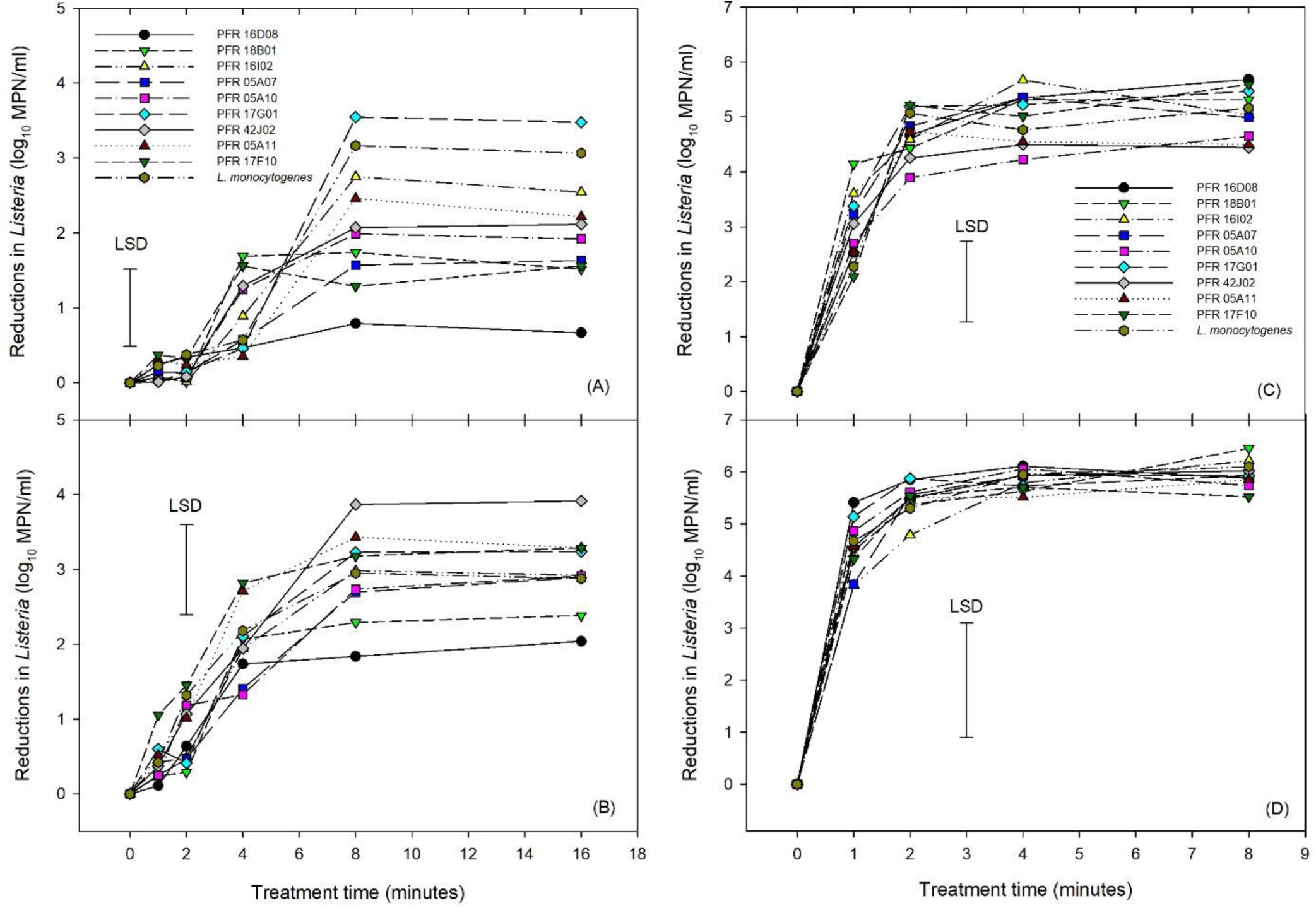
Reductions in numbers of *Listeria innocua* and of organisms in a *Listeria monocytogenes* cocktail during exposure to Tsunami™ 100 at 10 ppm (A), 20 ppm (B), 40 ppm (C) and 80 ppm (D). Error bars = least significant differences (*p* = 0.05, n = 2).

### Heat treatment

We employed mild heat as thermal inactivation procedure to assess the *Listeria* strains. Figure 4 represents the responses of different strains of *Listeria* to heat treatment which shows log_10_ reductions of *Listeria* during heat treatment at 55°C, over time up to 120 minutes. The *LM* cocktail showed log_10_-linear reductions up to 6 log_10_ MPN/mL at 60 min and then was totally inactivated by 120 min, as were all *L. innocua* strains. Different *L. innocua* strains had significantly different responses to the heat treatment. Strains PFR 17F10, PFR 05A07 and PFR 42J02 were all substantially more heat sensitive than the *LM* cocktail with total inactivation occurring within 30 min. PFR 17G01 also showed significantly more heat sensitivity than the *LM* cocktail after 15 and 30 min of heat treatment but by 60 min its inactivation rate was reducing, resulting in a total inactivation very similar to the *LM* cocktail. Similarly, none of the other *L. innocua* strains consistently matched the inactivation rates of the *LM* cocktail. While all showed similar reductions at 15 min, two strains (PFR 18B01 and PFR16I02) showed significantly more reduction at 30 min while the other three (PFR 05A10, PFR 05A11 and PFR 16D08) showed significantly less reduction at 60 min. The most similar was PFR 05A10 which had similar sensitivity to heat after 15 and 30 min and was only slightly more resistant than the *LM* cocktail after 60 min with a log_10_ reduction of 5.07 compared with a 6.06 log_10_ MPN/mL for the *LM* cocktail (LSD = 0.914 log_10_ MPN/mL).

**Figure 4.**
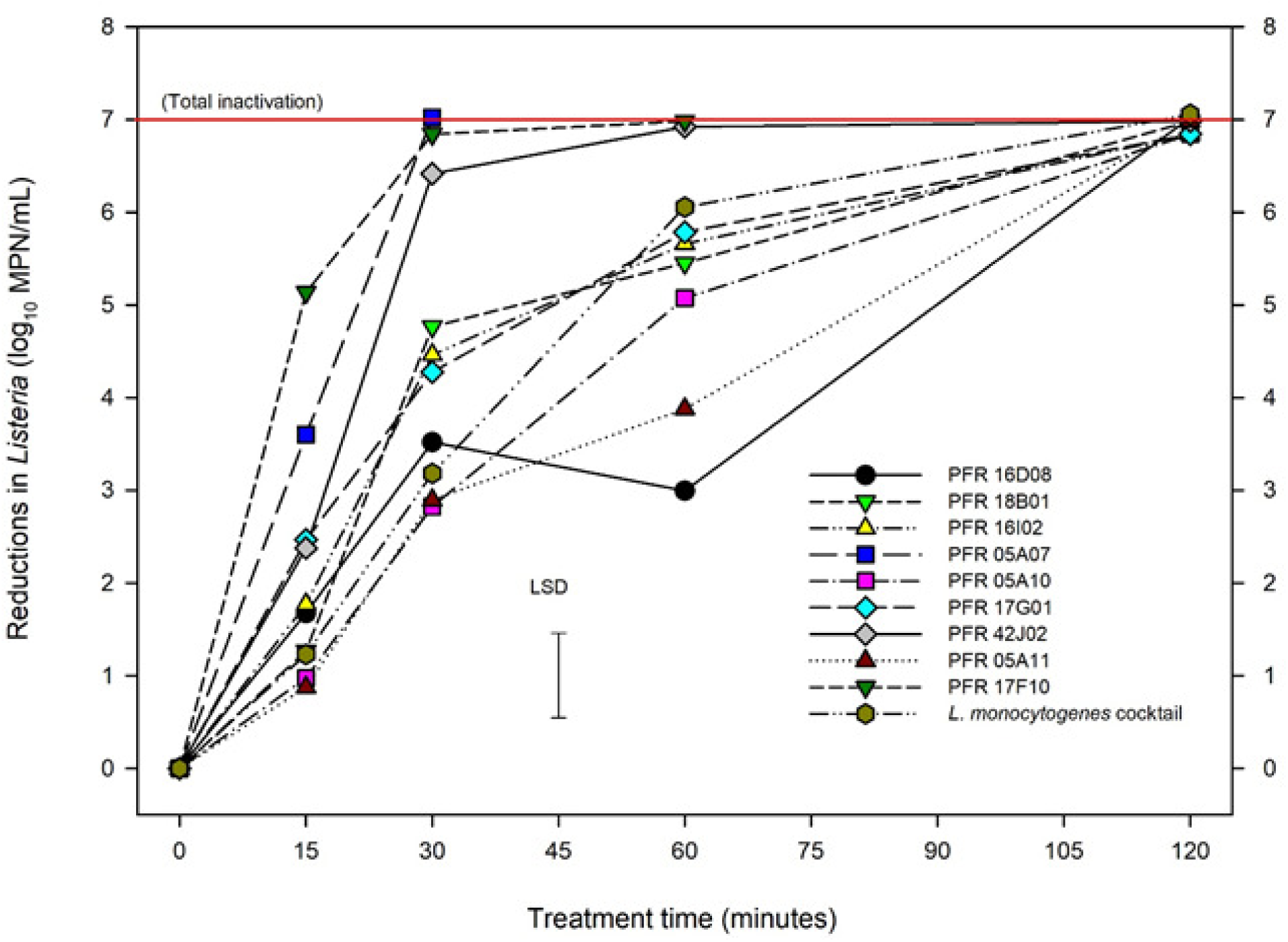
Reductions in numbers of *Listeria innocua* and of organisms in a *Listeria monocytogenes* cocktail at 55°C. Error bar = least significant difference (*p* = 0.05, n = 2).

### Hurdle treatments

For the combination treatment (hurdle), we conducted two hurdle treatments for *L. innocua:* one for individual strains (hurdle individual treatment) and the other for *L. innocua* cocktails (hurdle cocktail treatment) that were compared with the LM cocktails. Figure 5 shows the log_10_ reductions for the nine individual *L. innocua* strains and the *LM* cocktail tested against the hurdle treatments. Figure 6 shows the log_10_ reductions for different combinations of *L. innocua* cocktails compared to that of the *LM* cocktail in the hurdle treatments.

**Figure 5.**
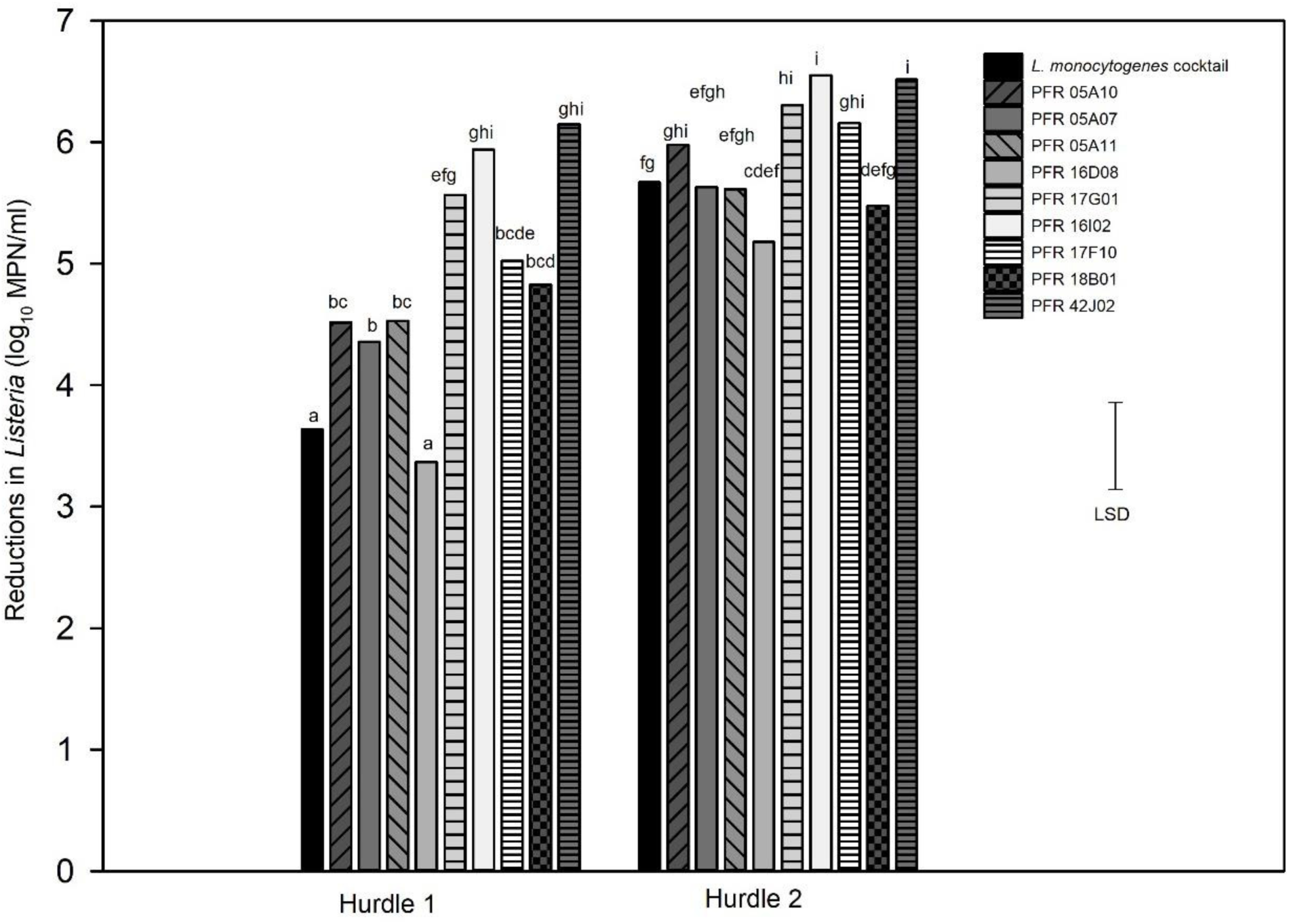
Hurdle individual treatments: Reductions in numbers in individual *Listeria innocua* strains and a *Listeria monocytogenes* cocktail in response to hurdle treatments: heat (55°C) for 7.5 minutes, UV-C at 328 mJ/cm^2^, Tsunami™ 100 at 10 ppm for 2 minutes (Hurdle 1) and heat (55°C) for 15 minutes, UV-C at 672 mJ/cm^2^, Tsunami™ 100 at 10 ppm for 4 minutes (Hurdle 2). Error bar = least significant difference (*p* = 0.05, n = 3).

**Figure 6.**
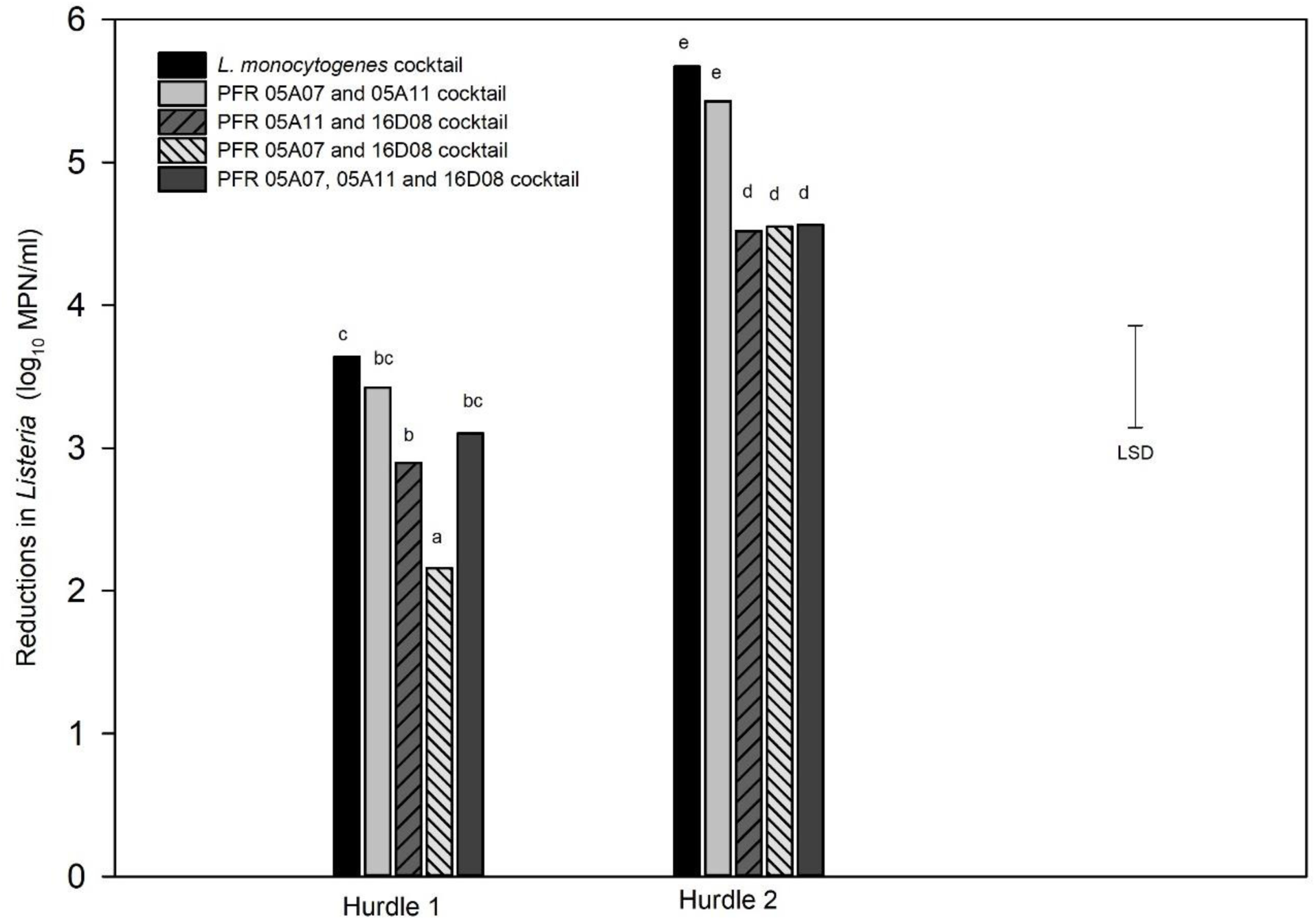
Hurdle cocktail treatments: Log_10_ MPN reductions in cell numbers of *Listeria innocua* cocktails and *L. monocytogenes* cocktail in response to hurdle treatments: heat (55°C) for 7.5 minutes, UV-C at 328 mJ/cm^2^, Tsunami™ 100 at 10 ppm 2 minutes (Hurdle 1) and heat (55°C) for 15 minutes, UV-C at 672 mJ/cm^2^, Tsunami™ 100 at 10 ppm 4 minutes (Hurdle 2). Error bar = least significant difference (*p* = 0.05, n = 3).

When tested individually, all but one strain (PFR 16D08) gave significantly higher log_10_ reductions in hurdle 1. In hurdle 2, PFR 05A07, 05A11 and 18B01 responded very similarly to the *LM* cocktail but only PFR 17G01, 16I02, 17F10 and 41J02 were significantly different, all being more sensitive to the hurdle treatments with higher reductions. Only PFR 16D08 did not differ significantly to the *LM* cocktail in its response to both hurdle combinations (Figure 5) but this strain was also substantially more resistant to all the treatments when applied individually (Figures 2, 3 and 4).

Cocktail combinations of three *L. innocua* strains (PFR 16D08, 05A07 and 05A11) that responded similarly to the *LM* cocktail were also subjected to the hurdle treatments (Figure 6). In both hurdle treatments the *L. innocua* cocktail containing PFR 05A07 and 05A11 was not significantly different to that of the *LM* cocktail. There was also no significant difference in the response of the cocktail of the three strains (PFR 05A07, 05A11 and 16D08) to hurdle 1 but in all other instances the *L. innocua* cocktail was significantly more resistant to the hurdle treatments than the *LM* cocktail.

*L. innocua* strains with similar responses to the *LM* cocktail are summarised in Table 1.

**Table 1.** Summary of the *Listeria innocua* strains that responded similarly to the *L. monocytogenes* cocktail under treatment conditions that differentiated the inactivation responses of *L. innocua* strains from each other. The strains were compared based on the least significant differences (*p* = 0.05,) for the individual or hurdle combination treatments.

| **Treatment** | **Treatment conditions** | ***Listeria innocua* strains showing similar responses (*P* <0.05) to those of the *LM* cocktail** |
| --- | --- | --- |
| **UV-C** | 1346 and 1009 mJ/cm<sup>2</sup> | PFR 05A10 and PFR 18B01 |
| **Tsunami™ 100 (10 ppm)** | 4 min | All except PFR 18B01 |
| **Tsunami™ 100 (10 ppm)** | 8 and 16 min | PFR 17G01, PFR 16I02, PFR 05A11, PFR 42J02, PFR 05A10 |
| **Tsunami™ 100 (40 ppm)** | 2 min | All except PFR 17F10 |
| **Heat (55°C)** | 15 min | All except PFR 17F10 and 05A07 |
| **Heat (55°C)** | 30 min | PFR 05A11, PFR 05A10, PFR 16D08 |
| **Heat (55°C)** | 60 min | PFR 17G01, PFR 16I02, PFR 18B01 |
| **Hurdle 1** | 55°C (7.5 min), 10 ppm Tsunami™ 100 (2 minutes) and UV-C (328 mJ/cm<sup>2</sup>) | PFR 16D08, cocktail of PFR 05A07 and PFR 05A11, cocktail of PFR 16D08, PFR 05A07 and PFR 05A11 |
| **Hurdle 2** | 55°C (15 min), 10 ppm Tsunami™ 100 (4 min), | PFR 05A07, PFR 05A11, PFR 05A10, PFR 18B01, PFR 16D08, PFR 17F10, cocktail of PFR 05A07 |
|  | UV-C (328 mJ/cm<sup>2</sup>) | and PFR 05A11 |

### Discussion

The aim of the present study was to select suitable strain(s) of *L. innocua* to be used as non-pathogenic surrogates for *L. monocytogenes* in laboratory and pilot-scale process studies. This meant identifying a surrogate whose responses matched those of an *LM* cocktail as closely as possible. Some of the criteria we considered in selecting a suitable surrogate were: a surrogate being more sensitive than LM could result in mild treatments being promoted which would not assure food safety, while being more resistant could result in selecting a harsh treatment that may have higher costs and could cause more damage to the product than necessary and, if no surrogate was found to totally match the pathogen then our aim was to choose one that was slightly more resistant so that food safety would be assured. We carried out independent experiments with 10 *L. innocua* strains and a cocktail of 11 *L. monocytogenes* strains isolated from New Zealand fresh produce using treatments suitable for fruit or vegetables that were to be eaten raw: UV-C, sanitiser, and heat (Figures 2 to 4).

The germicidal effect of UV-C irradiation has long been known and has been employed widely to inactivate indicator organisms and pathogens in fresh produce, environmental biofilms, food products and ready-to-eat food products (27, 29, 42–48). Montgomery and Banerjee used pulsed UV light (PUV) for 10 and 20 seconds to treat *L. monocytogenes* and *E. coli* biofilms on the surfaces of lettuces. It was observed that longer PUV exposure time and shorter light source distance to the sample (20 s—4.5 cm) resulted in a significant viable cell reduction of both pathogens compared to shorter exposure time and longer light source distance (10 s—8.8 cm) (31). Similarly, in our study we observed increases in reduction rates with increased exposure time and shorter light source distance. On average all our *L. innocua* strains were more resistant to UV-C than the cocktail of 11 diverse strains of *L. monocytogenes* with all strains except PFR 18B01 and PFR 05A10 being significantly more resistant at doses of over 1000 mJ/cm^2^. This suggests that in general *L. innocua* is more resistant to UV-C than *L. monocytogenes*. Previous studies have used UV-C to inactivate *L. monocytogenes* from meat, processed food products and fresh produce (27, 43, 45, 46, 48–50). However, these studies cannot be compared directly to ours due to the differences in matrices, dose of UV-C, and experimental procedures.

Similarly, in the food industry, sanitisers and cleaning agents have been used in cleaning regimes for decades to reduce microbial contamination, thereby improving product shelf life and food safety. Previous studies have investigated the inactivation efficiency of Tsunami™ 100 (acetic acid, peroxyacetic acid and hydrogen peroxide) in different fresh produce (5, 36, 51–53). These studies indicated that peroxyacetic acid-based sanitisers are effective at lower concentrations compared to other sanitisers and were effective at killing *Listeria* spp. and their biofilms. In this study, as expected, we observed that the inactivation efficiency of Tsunami™ 100 increased with increased concentration and time of exposure. The United States Environmental Protection Agency (US EPA) has recommended 30–80 ppm for 45 seconds to yield an effective inactivation of non-pathogenic spoilage organisms on fresh produce and 30 minutes for soil pathogens (54). When applying Tsunami™ 100 to *L. innocua* cultures suspended in clean water, we observed rapid reductions in numbers at 40 and 80 ppm. Maximum inactivation (4 and 5 log_10_ reductions at 40 ppm and 80 ppm, respectively) was not observed until exposure times exceed 2 and 1 min, respectively (Figures 3C and 3D). The lower concentrations of 10 and 20 ppm also gave increasing inactivation of most *L. innocua* strains with increasing contact time up to 8 minutes (maximum reductions all below 4 log_10_) after which further extension of contact time did not show any additional benefit (Figures 3A and 3B). In some of the more resistant strains, activation stopped after just 4 min (PFR 16D08 and PFR 18B01 at both 10 and 20 ppm and PFR 17F10 just at 10 ppm). At the lowest tested concentration of 10 ppm, just two *L. innocua* strains (PFR 16I02 and PFR 17G01) had very similar log_10_ reductions in numbers to that of the *LM* cocktail while the other strains were more resistant to sanitisers than the *LM* cocktail.

In the heat treatment, the survival rate decreased with extended heating time, which was expected and has been observed by other researchers (55). All strains had been totally inactivated after 120 minutes at the relatively mild temperature of 55°C. Such an exposure time is unlikely to be practical as a post-harvest treatment and different strains responded differently to heat treatment after shorter exposure times.

In another thermal inactivation study evaluating *L. innocua* strains as surrogates, E. C. Friedly et al. (25) used five different *L. innocua* strains M1, 5639, 5640, and 2745 (from Special *Listeria* Culture Collection, Univ. of Wurzburg, Germany) and tested them against the higher temperatures (62.5 to 70°C) suitable for hamburger patties. They recommended using M1 as a surrogate as it had the “greatest” margin of safety. M1 is well characterised and has been used by other researchers for *Listeria* surrogate work around the world (13, 25) and this strain has been used in thermal inactivation studies of various products in the laboratory environment (25). However, the temperatures used by E. C. Friedly et al. (25) are too high for fresh produce due to its sensitivity to heat. The approach of using the organism with the greatest margin of safety (i.e. the most resistant) as a surrogate would be likely to lead to using processes that give unacceptable losses in quality much milder than those applied to milk and meat.

In our lab-based study all *L. innocua* strains had significantly different (*p* < 0.05) responses to the *LM* cocktail in one or more of these individual treatments. Across the three treatments, PFR 05A10 was the most similar in its responses compared to the *LM* cocktail, only being significantly different at 55°C for 60 min (Table 1). This strain invariably gave responses that showed it to be either very similar or slightly more resistant to the individual treatments than the *LM* cocktail (lines below that of the cocktail in Figures 2–4). If a single strain were to be used to evaluate all the single treatments, this strain would be the best surrogate of those tested. However, for the sanitiser, although PFR 05A10 was not significantly different to the *LM* Cocktail, PFR 16I10 was consistently more similar to the *LM* cocktail than PFR 05A10 so this strain might be a better surrogate for studies on peroxyacetic acid-based sanitisers like Tsunami™ 100. Like PFR 05A10, it was also slightly more resistant than the *LM* cocktail.

Given that *L. monocytogenes* can survive under harsh conditions and only mild treatments can be applied to fresh produce, inactivation of *L. monocytogenes* using a single treatment is unlikely to provide a post-harvest regime sufficient to achieve desired reductions in bacterial numbers. The concept of combined application of pathogen inactivation procedures has been accepted and widely used for the past decade. In our study we found that in each individual treatment (UV-C, Tsunami™ 100 and heat) gave a different profile of *L. innocua* strains that responded most closely to the *LM* cocktail. This suggests that in a real-world scenario, each *L. innocua* strain will respond to UV-C, sanitiser and heat differently and a cocktail of surrogates would be appropriate for use in inactivation studies. For example, strains PFR 18B01 and PFR 05A10 responded most similarly to UV-C exposure, as did the *LM* cocktail, whereas, PFR 16I02 and PFR 17G01 responded most similarly to the sanitiser treatment at the lower concentration of 10 ppm and the strain PFR 05A10 was most similar in the heat treatment.

When we combined the three treatments into a series of hurdles, dramatically greater reductions in numbers were achieved for the individual strains (Figure 5) compared to when the single treatments were applied (Figures 2–4). In the milder hurdle combination (Hurdle 1) that was applied at about half the strength as in Hurdle 2, all strains suffered more than 3 log_10_ reductions with some reaching 6 log_10_ reductions. The UV-C, sanitiser and heat treatments used in the Hurdles did not achieve significant reductions for any of the strains when applied as individual treatments but when combined as a hurdle combination, all strains suffered at least 5 log_10_ reductions in numbers. This demonstrated that these three treatments work synergistically. In general, the synergistic hurdle effect of the three combined treatments was mirrored in both the individual strains and the cocktails. Although hurdle technology has been applied to different fresh-cut-produce and food products (41, 56, 57), our laboratory-based hurdle study cannot be compared with these studies as they involved different treatments as combinations including spray washing, essential oils, high-pressure processing, sonication and other as suited for different food products.

Several strains including PFR 16D08 responded to Hurdle 2 in a similar manner to the *LM* cocktail but only PFR 16D08 was as resistant to the mild Hurdle 1 treatment as the *LM* cocktail. PFR 05A10 that responded similarly to the *LM* cocktail when the treatments were applied individually was significantly more sensitive when they were applied as a combination in Hurdle 1. If such a strain were to be used as a surrogate, it could result in researchers underestimating the effect of a treatment against *L. monocytogenes* and potentially leading to sale of unsafe product. While PFR 16D08 on its own could be selected as a surrogate for this combination of treatments, when the treatments were applied individually, PFR 16D08 was usually much more resistant than the *LM* cocktail. This was of concern as, when applied for studies on actual produce, it might be found necessary to apply some of the hurdles at even lower intensities than were applied in Hurdle 1. For example, some produce might be damaged at 55°C and holding product in such a treatment for 7.5 min might be too long for high-throughput industries. If the intensities of one or more of the treatments had to be reduced the combination system might respond more similarly to the single treatment. In this case, if PFR 16D08 were being used, it is likely that more intense treatments would be adopted than necessary to achieve target reductions in *L. monocytogenes*, and there is a danger of recommending treatment regimes, that may not be suitable for sensitive or highly perishable fresh produce, causing unacceptable damage to the produce and costing the industry more. Higher sanitiser concentrations might also be recommended, leading to unnecessary chemical residues.

We also tested the responses of three cocktail combinations of *L. innocua* to the two hurdle treatments (Figure 6). As with the individual treatments, the cocktails that included PFR 16D08 were consistently more resistant to the hurdle treatments than the *LM* cocktail. However, the responses of the cocktail of PFR 05A07 and PFR 05A11 were not significantly different to those of the *LM* cocktail. When challenged with Hurdle 2 as individual strains, PFR 05A07 and PFR 05A11 responded almost exactly the same as the *LM* cocktail although they were more sensitive to the Hurdle 2 combination (Figure 5). Both were more resistant to UV than the *LM* cocktail (Figure 2) so have the same potential limitations as PFR 16D08 in this regard. They were more similar in their response to the sanitiser (Figure 3) than PFR 16D08 and, although PFR 05A10 was very sensitive to heat inactivation, this was balanced by PFR 05A11 which was more similar in its response than PFR 16D08, particularly after 60 min exposure. Overall, from the data presented in this paper, when investigating individual and hurdle combinations to inactivate *L. monocytogenes*, we recommend using not an individual as surrogate but the cocktail of PFR 05A08 and PFR 05A11 although others might prefer to use just PFR16D08

To conclude, nine genetically dissimilar *L. innocua* strains and 11 *L. monocytogenes* strains (combined in a cocktail) were compared by treating them with UV-C, sanitiser and heat, and a combination of these three in an effort to select suitable surrogate candidate(s). The results indicated that each *L. innocua* strain responded differently. The hurdle treatment produced a synergistic inactivation that had a significant reduction in the survival rates for the individual species and the cocktails. Our study indicated that a cocktail of PFR 5A08 and PFR 5A11 strains may serve as a good surrogate for fresh produce thermal, UV-C, sanitiser and hurdle studies with a significant safety margin. Testing these strains in different food matrices and post-harvest hurdle treatment regimes, will provide insights into recommended heating times for inactivating *Listeria* spp. and/or *L. monocytogenes* in food products and recommended dose-time combinations for inactivating *Listeria* in fresh produce. The study highlights that it is important to test more than one surrogate strain to obtain an effective inactivation regime in food products as different strains exhibit different responses to inactivation procedures. The current study was carried out in laboratory media, but future studies should test different food matrices for validating the *L. innocua* strains to be used as potential surrogates.

## Materials and methods

### Bacterial cultures

Based on their Pulse Field Gel Electrophoresis (PFGE) pulsotypes, genetically diverse *Listeria* strains were selected from The New Zealand Institute of Plant & Food Research Ltd (PFR) Culture Collection. Pure bacterial cultures were revived from −80°C in tryptic soy broth plus 0.6% yeast extract (TSBYE, Bacto™, BD, Spark, USA). Cultures were plated onto TSAYE agar (TSBYE plus 1.5% agar (Bacto) and incubated for 48 hours at 37°C. From these plates, 3 mm diameter colonies were selected (measured using a digital vernier calliper (Model 071701, ROK International Industry, China)) and inoculated into TSBYE. Cultures were incubated for 48 hours at 37°C allowing the cultures to reach the stationary phase. *L. innocua* strains from diverse New Zealand sources (n=19) were typed using *Asc*I. The restriction patterns were analysed using the InfoQuest™FP molecular analytical software tool (Bio-Rad, USA) and the similarity indices were used to build the dendrogram (Figure 1A) to analyse the genetic similarity and/or dissimilarity. Good resolution of the *L. innocua* isolates was found with just one enzyme. To conduct the lab-based pilot study, nine genetically diverse (genetic dissimilarity of at least 35% on the dendrogram) *L. innocua* strains (PFR 05A07, PFR 05A10, PFR 05A11, PFR 16D08, PFR 16I02, PFR 17F10, PFR 17G01, PFR 18B01 and PFR 42J02) were selected. *L. monocytogenes* strains from horticultural sources (n=10 were characterised by PFGE, using two rare cutting restriction enzymes, *Asc*I and *Apa*I, and analysed using the InfoQuest software. Ten *L. monocytogenes* strains (Figure 1B) were selected based on their genetic diversity and on the horticultural sources that they were isolated from (PFR 41E01, PFR 41E03, PFR 41E05, PFR 41F08, PFR 41H07, PFR 40I05, PFR 40I07, PFR 41J05, PFR 41J08, and PFR 41J09). The *L. innocua* strains were individually compared to a cocktail of the 10 stains of *L. monocytogenes* plus an ATCC reference *L. monocytogenes* strain (Scott A, PFR 16B03) (*LM* cocktail). The *LM* cocktail was prepared by mixing 1 mL of stationary phase culture from each of the *L. monocytogenes* strains. Counts of the cultures after the 48-hour incubation period averaged 6.5 log_10_ MPN/mL and the OD of the *LM* cocktail was between 0.5–0.6 measured on PD-303 spectrophotometer (Apel, Co. Ltd, Japan). Each experiment assessed the effect of the inactivation treatments against each of the 10 *L. innocua* strains and the *LM* cocktail.

### *Listeria* quantification

All bacterial counts were enumerated using the MPN technique for *Listeria* according to the method of the FDA Bacterial Analytical Manual (58, 59). Cultures were serially diluted in triplicate in 96 well plates up to 10^−15^. These plates were incubated for 48 hours at 30°C after which aliquots of 2 µL were plated onto ChromAgar (750006, Paris) *Listeria*. Plates with typical growth characteristics (green colonies for *L. innocua* and green with a halo for the *LM* cocktail) were recorded for enumeration using the FDA MPN method spreadsheet (59).

### UV treatment

The UV-C treatment was conducted in the biosafety level II cabinet equipped with two lamps, a Philips UV lamp (TUV 30 watt, G30 T8, Philips, The Netherlands) and a Sankyo Denki UV germicidal lamp (G15T8, Sankyo Denki, Japan) installed side by side in the cabinet. UV-C irradiance measurements (µW/cm^2^) were carried out using a UM-10 Konica Minolta UV radiometer. To obtain different doses of UV-C light, exposure plates of cultures were placed at different distances (55 cm, 45 cm, 34 cm and 26 cm) from the UV-C lamp for 20 min. To calculate UV-C dose, the radiometer was left in the cabinet at the respective distances and UV-C irradiance (µW/cm^2^) was recorded at 15 seconds intervals. UV-C doses after 20 minutes’ exposure at the respective distances from the lamp were 600, 672, 1009 and 1346 mJ/cm^2^. For one of the hurdle treatments described below, cultures were placed 45 cm from the lamp for just 10 minutes which gave a dose of 328 mJ/cm^2^.

Ten microliters of 48-hour *L. innocua* cultures or the *LM* cocktail were dispensed into individual wells in 12 well plates (Corning-Costar, CLS3512, supplied by Sigma-Aldrich). The plates were exposed to UV-C light for 20 minutes. Immediately after UV-C treatment, 490 µL of Buffered *Listeria* Enrichment Broth (BLEB, Acumedia, USA) was added into each well, and thoroughly mixed by repeated pipette aspiration. These suspensions were transferred into a sterile 96 well plate (Greiner Bio-One, GmbH, Frickenhausen, Germany) and survivors quantified using the MPN method.

### Sanitiser treatment

Tsunami™ 100 (Ecolab Inc., Minnesota, USA) was diluted to working concentrations of 10, 20, 40 and 80 ppm, and the concentration was confirmed as peroxide equivalents with a Palintest Photometer 5000 using the hydrogen peroxide HR test at 490nm. The cultures were exposed to the sanitiser for 1, 2, 4, 8- and 16-minutes following procedures adopted from Cruz and Fletcher (5). One mL of each culture was centrifuged at 15700 RCF for 5 minutes. The supernatant was removed, and cells washed with sterile water and re-suspended in 1 mL sterile water. Cell suspensions (50 µL) were dispensed into a 96 well plate and 50 µL of the sanitiser was added into each well. Plates were then left at room temperature for the designated contact time. To terminate the sanitiser reaction, 150 µL of neutraliser solution (containing 5% egg yolk emulsion (Difco), 1% sodium thiosulphate (AnalaR, BDH, Chemicals Ltd, Poole, England) and 0.5% Tween® 80 (Spectrum, Gardena, CA) in TSBYE was added to each well containing the bacterial culture and sanitiser. Neutralised suspensions were then serially diluted in BLEB up to 10^−15^, and survivors quantified using the MPN method.

### Heat treatment

All *L. innocua* strains and the *LM* cocktail were exposed to 55°C for 15, 30, 60, 120, 240, and 480 minutes to investigate the inactivation of the isolates. An aluminium heating block (model D1105, Labnet Intl. Inc. Edison, NJ, USA) was placed in a water bath (type TC120, Grant Instrument (Cambridge Ltd), Shepreth SG8 6GB, England). Capped micro-tubes (0.6ml thin walled Axygen™ MaxyClear Snaplock Microtubes, type MCT060R, Axygen Scientific Inc. NY, USA) containing 490 µL of BLEB were placed in the heating block which was then heated up to 55°C and left for 30 minutes to equilibrate to the test temperature before the cultures were added. Capped micro-tubes fitted with calibrated type-T thermocouples were used to measure the temperature and the temperature was measured at 1 sec interval using a 1000 series Grant Squirrel meter/logger (Grant Instruments Ltd, Cambridge, England). When the media had equilibrated at the desired test temperature, 10 µL of 48-hour 30°C cultures were added to each micro-tube and mixed three times by aspiration using an automated pipette. There was a temperature drop (average = 2.64°C) while adding the cultures to the heated media and on average it took 2.23 minutes to return to the test temperature.

Bacterial counts were taken at T0 – Untreated Control and at Ts – specified time intervals at the test temperature. At each sampling interval microtubes were placed on ice and were then serially diluted in BLEB in 96 well plates. Surviving cells were enumerated by the MPN method.

### Hurdle treatments

All *L. innocua* strains used in the present study were tested individually in the hurdle treatments. We conducted two hurdle treatments for the *L. innocua* individual strains and cocktails that were compared with the LM cocktails. We refer to these two hurdle treatments as ‘hurdle individual treatment’ and ‘hurdle cocktail treatment’. In both hurdle treatments, hurdle 1 consisted of a combination of milder treatments, sequentially heat (55°C for 7.5 minutes) followed by UV-C (328 mJ/cm^2^) and sanitiser (10 ppm of Tsunami™ 100 for 2 minutes). Hurdle 2 treatment (harsh treatment combinations) consisted of a combination of heat (55°C for 15 minutes) followed by UV-C (672 mJ/cm^2^) and sanitiser (10 ppm of Tsunami™ 100 for 4 minutes). The procedures for each hurdle treatment were identical to the independent treatment procedures. The samples were heat-treated in capped micro-tubes, transferred into 6 well tissue culture plates for UV-C exposure, then transferred to a 96 well plate for sanitiser treatment and finally neutralised. The samples were then serially diluted and the survivors were enumerated using the MPN method.

For the hurdle cocktail treatment, cocktails of *L. innocua* and LM cocktail were compared. The *L. innocua* strains used in the cocktails were selected based on their individual responses in comparison to the *LM* cocktail in the individual treatments conducted previously. The cocktails selected were (1) PFR 05A07 and 05A11; (2) PFR 05A11 and 16D08; (3) PFR 05A07 and 16D08; and (4) PFR 05A07, 05A11 and 16D08. These *L. innocua* cocktails were subjected to the hurdle 1 and 2 treatments as described above.

### Statistical analysis

Each independent experiment was carried out twice while the hurdle combination experiments were carried out three times, each using each individual strain of *L. innocua* and the *LM* cocktail. The calculated MPNs were log-transformed, log_10_ reductions were calculated and the variance was stabilised. The standard errors of the mean were calculated and one-way analysis of variance (ANOVA) (Genstat Version 17, 2016) and a post-hoc Fisher’s Least significant differences (LSD, *p* = 0.05) were calculated. These parameters were used to compare individual *L. innocua* strains, the *LM* cocktail and *L. innocua* cocktails and were presented graphically as LSD bars.

## Acknowledgements

The research was funded through the Ministry of Business, Innovation & Employment (MBIE) and New Zealand Apples & Pears Incorporated (NZAPI) Partnership Programme PNZEA1401. We thank our Biometrician, Mr Duncan Hedderley for the statistical assistance.

